# Super-elasticity of plasma- and synthetic membranes by coupling of membrane asymmetry and liquid-liquid phase separation

**DOI:** 10.1101/2020.07.13.198333

**Authors:** Jan Steinkühler, Tripta Bhatia, Ziliang Zhao, Reinhard Lipowsky, Rumiana Dimova

**Affiliations:** Theory and Bio-Systems, Max Planck Institut of Colloids and Interfaces, Science Park Golm, 14424 Potsdam, Germany

## Abstract

Biological cells are contained by a fluid lipid bilayer (plasma membrane, PM) that allows for large deformations, often exceeding 50% of the initial (or projected) PM area. Biochemically isolated lipids self-organize into membranes, but the extraordinary deformability of the plasma membrane is lost. Pure lipid bilayers are prone to rupture at small (<2-4%) area strains and this limits progress for synthetic reconstitution of cellular features such as migration, phagocytosis and division. Here, we show that by preserving PM structure and composition during isolation from cells, vesicles with cell-like elasticity are obtained. We found that these plasma membrane vesicles store significant area in the form of nanotubes in their lumen. These are recruited by mechanical tension applied to the outer vesicle membrane showing an apparent elastic response. This “super-elastic” response emerges from the interplay of lipid liquid-liquid phase separation and membrane asymmetry. This finding allows for bottom-up engineering of synthetic vesicles that appear over one magnitude softer and with three fold larger deformability than conventional lipid vesicles.

---

After chemical cleavage of the cytoskeleton – PM anchors, turgor pressure and membrane tension lead to the formation of spherical blebs, called giant plasma membrane vesicles (GPMVs). These vesicles reconstitute fluid PM lipid and membrane protein^1^, including soluble cytosolic components (**Fig. 1a** and Methods)^2,3^. GPMVs were isolated from the cells and stained with the membrane dye FAST-Dil (see Methods for details). Unexpectedly, fluorescence intensity was observed not only on the outer GPMV membrane, but appeared also in the vesicle lumen (**Fig. 1b**). Deconvolution and super resolution imaging (STED) revealed a tubular lipid network which is contained within the isolated GPMVs and connected to the outer membrane (**Fig. 1c**). In some vesicles, the tubular network was found to be very dense and possibly branched (**Fig. 1c insert**). The diameter of the lipid nanotubes appeared to be mostly below the resolution limit of our STED setup (about 60 nm), but some tubes were wide enough to be directly resolved by STED measurements (**Fig. 1d**). To aid interpretation of the imaging results, a water soluble dye was added to the outer aqueous solution. Fluorescence signal from the water channels formed by the nanotubes provided direct evidence for the tubular structure and connection to the outer membrane (sulforhodamine B dye in **Fig. 1e**). Together, this confirms that GPMV isolated in our conditions reconstitute a nanotubular lipid network (**Fig. 1f**) which is similar to tubular structures found in cells^4,5^. Curvature elasticity arguments suggest that, coexistence of the outer, weakly curved membrane segment, and the highly curved nanotubes requires that the membrane exhibits preferred, or spontaneous curvature, in the order of m_tubes_> 1/100 nm^−1^ = 10 μm^−1^, stabilising the highly bent lipid bilayer^6^. Similar spontaneous curvature was shown to be generated by adsorption of curvature-inducing BAR domain protein to the bilayer^7^. Indeed proteomics indicate that BAR-domain proteins are present in GPMVs ^3^ and by equilibration of the binding free energy these should be found primarily bound to the nanotubes ^8^.

**Figure 1.**
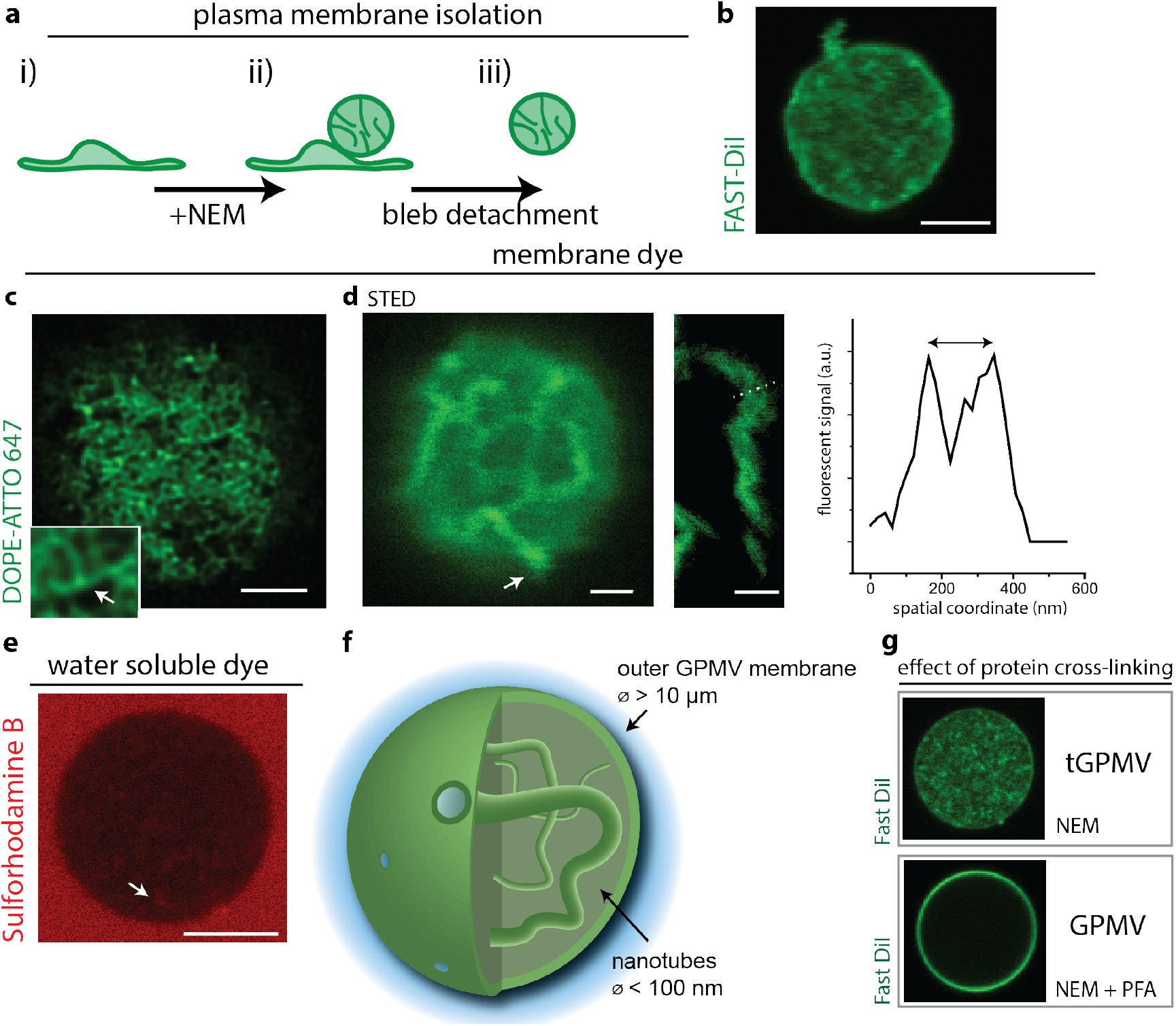
**a)** Extraction and isolation of plasma membrane by incubation of U2OS cells by incubation with N-Ethylmaleimide (NEM). **b)** Membrane extracts were isolated and labelled with fluorescent membrane dye (FAST-DiI) (shown in green). **c)** High-resolution imaging of the internal structures by deconvolution confocal imaging reveals dense lipid network. Possible 3-way branches in the network are shown in the insert. **d)** Stimulated emission depletion (STED) microscopy of internal structures found in the lumen of the vesicles. Nanotubes appear to be connected to the outer membrane. Scale bar 1 μm. Right insert: Two peaks in fluorescence intensity plot along the dashed line indicated on the image. The two peaks correspond to the nanotube walls and exemplify a nanotube radius of about 100 nm. Scale bar 500 nm. **e)** Diffusion of water-soluble dye added to the outer solution into the nanotubes. Arrow shows signal stemming from the water-filled nanotube interior. **f)** Sketch of the overall GPMV structure (vesicle and nanotube diameter not to scale). **g)** Effect of crosslinking chemical PFA on plasma membrane extracts. We term tGPMV extracts those that reconstitute a tubular lipid network. If not indicated otherwise all scale bars 5 μm.

Consistent with protein coverage on the cytosolic leaflet of the nanotubes, addition of the protein crosslinker paraformaldehyde (PFA) to the vesicle lumen led to crosslinking of the nanotubes. This effect was exploited to obtain the high-resolution images of the otherwise dynamically fluctuating nanotube network shown in **Fig. 1c,d** (also see Methods).

However, if the crosslinking chemicals were present *during* plasma membrane extraction, the GPMVs interior appeared free from any fluorescence signal of membrane dye (**Fig. 1g** and Methods), suggesting that the nanotubes network was retrained inside the cells by the crosslinking agent. Consistent with previous reports^9^, sometimes the membranes isolated in cross-linking conditions appear to be permeable to the water-soluble dyes. However this effect leads to a homogenous dye distribution in the vesicle lumen and can be clearly distinguished from the structured signal from internal nanotubes shown in **Fig. 1e**. Previous experiments using GPMVs were often conducted in the presence of crosslinkers^10^, and to differentiate we term our vesicles reconstituting a tubular lipid network **tGPMVs**.

## Tension retracts nanotube network to outer membrane

As long as all membranes remain fluid and the nanotubes in the vesicle lumen are connected to the outer membrane, it should be possible to retract them to the outer vesicle membrane by applying membrane tension. Indeed, when tGPMVs were aspirated into glass capillaries, similar to those used for elastic probing of cells^11^ and lipid vesicles^12^, we were able to induce large-scale elastic deformations corresponding to about 40% increase of the initial (spherical) apparent outer membrane area **(Fig. 2a)**. Along the aspiration trajectory tGPMVs remained stably aspirated at a given tension and without any indication of hysteresis (**Fig. 2a)**. GPMVs that were isolated in conditions that do not reconstitute a nanotube network, appeared about 3 orders of magnitude stiffer and rupture at small area strains of about 3%, reminiscent of pure lipid bilayers^12^(**Fig. 2b**). When compared to literature data of micropipette aspiration of neutrophil cells^11^, it becomes clear that tGPMV elasticity appears similar to intact cells. While elasticity and area regulation in cells is certainly orchestrated by a range of (active) processes, including the actomyosin cortex and exocytosis of vesicles, our results suggest, that the plasma membrane itself is positioned to exhibit an elastic response matching the cortical tension. Next, we aimed to understand the essential features that enable PM super-elasticity.

**Figure 2.**
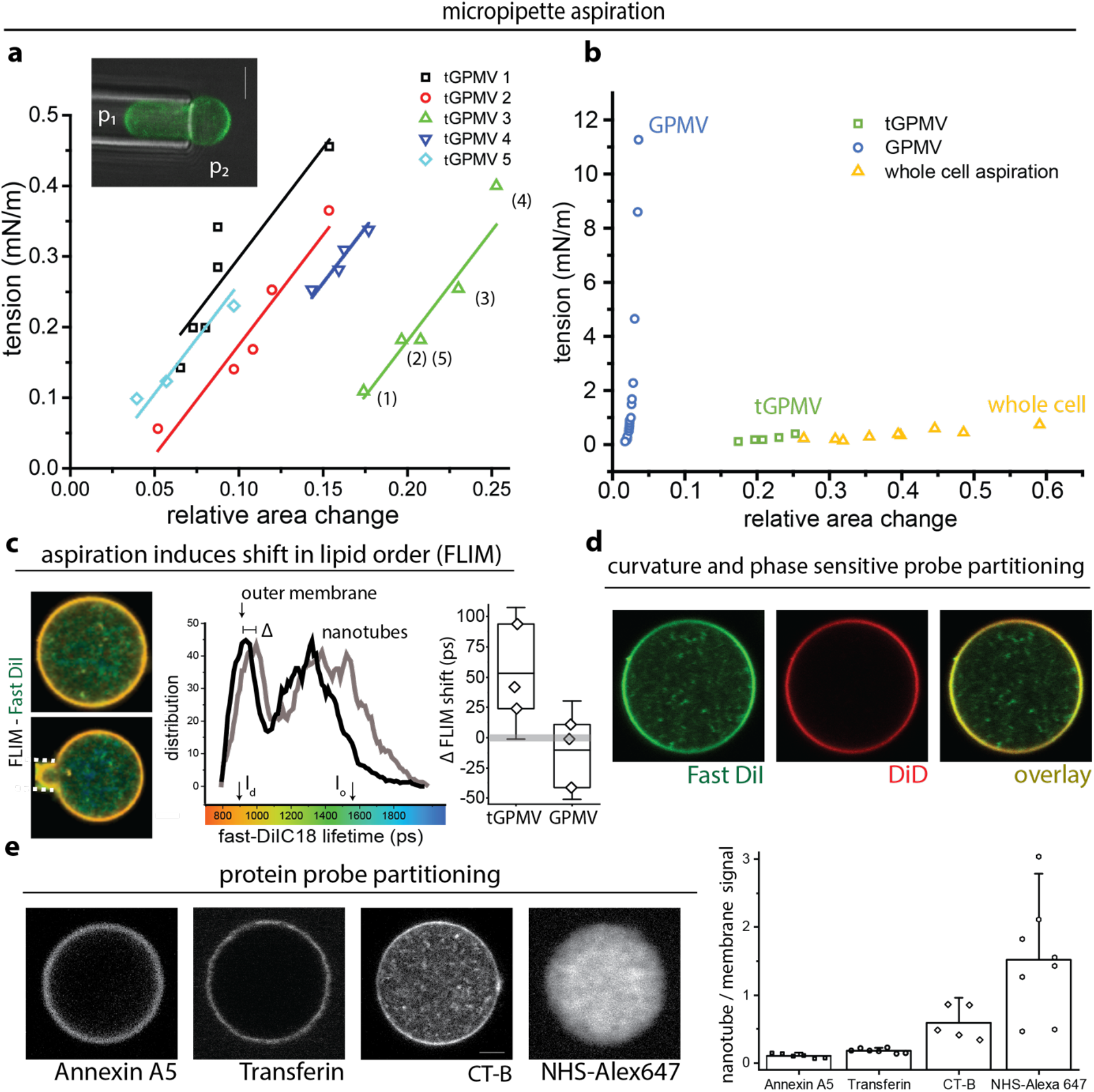
**a)** Micropipette aspiration of individual tGPMVs (colour-coded) at varying pressure difference Δp=p_2_-p_1_, the tension is calculated from the Laplace pressure 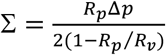, where *R_p_* and *R_v_* are the corresponding pipette and spherical membrane segment radii. The numbers in brackets indicate the experimental sequence of applied pressures. **b)** Comparison between apparent elastic moduli obtained for GPMVs, tGPMVs and neutrophils (data from^11^). **c)** FLIM data, colour code indicates FAST-DiI fluorescence lifetime in a free tGPMV and when aspirated (bottom image). White lines indicate micropipette. Histogram shows lifetime distribution (black – free tGPMV, grey – aspirated tGPMV). Upon application of a small membrane tension (0.7 ± 0.3 mN/m), the lifetime distribution of the outer membrane is shifted to higher values as indicated in the grey trace. We quantify the tension-induced change (Δ) in peak lifetime of the outer segment of tGPMVs and GPMVs. Each datapoint represents an independent experiment. **d)** Co-staining of the same tGPMV with membrane dyes FAST-Dil and DiD (green and red channel) and overlay. **e)** Staining of tGPMV with protein (amine groups) binding NHS-Alex647 and fluorescently conjugated Annexin A5, Transferrin and Cholera toxin subunit B (CT-B) (see Methods for details) and signal ratio between protein and membrane dye in the outer membrane of tGPMVs. All vesicle diameters were between 10 μm and 30 μm.

## Nanotube retraction leads to phase change of the outer membrane segment

It seems likely that the extraordinary tGPMV deformability proceeds by the recruitment of the nanotubes into the outer membrane, similarly to behaviour of tubulated lipid vesicles^13,14^. Indeed, recruitment of membrane materials from the nanotube could be directly visualized by fluorescence lifetime measurements (FLIM): FAST-DiI FLIM is a measure of membrane composition and is related to the order parameters of the acyl chain over a wide range of membrane composition^15^. Colour coding for FAST-DiI FLIM immediately reveals two distinct populations of fluorescence lifetimes between the outer membrane segments and nanotubes (**Fig. 2c**). Indicating a stark contrast in composition and lipid order between the outer membrane segment and the nanotubes.

Next, we observed the fluorescence lifetime of the outer membrane segment upon aspiration of a tGPMV. We found that the fluorescence lifetime of the dye in the outer membrane segment shifts by Δ towards to the lifetime found in the nanotubes (**Fig. 2c**). Control aspiration experiments on GPMV (**Fig. 2c**) exhibit no such shift in lifetime (Δ≈0). Thus, these measurements represent direct evidence of mixing of the two membrane segments by applying tension to the outer membrane segment. Essentially, we whiteness a continuous phase-state change in the outer membrane segment with increase in aspiration pressure and membrane tension.

## Phase separation of outer membrane segment and nanotubes lead to distinct membrane compositions

The difference in composition between nanotubes and outer membrane segment will be essential for the interpretation of the super elastic response and was corroborated by further experiments. Co-staining with the dyes FAST-DiI and DiD reveals segregation of the two molecules between the outer membrane segment and the highly curved nanotubes (**Fig. 2d**). FAST-DiI and DiD have similar molecular structure, except that FAST-DiI exhibits a double bond in the acyl chain. Compared to this small chemical difference, a strikingly different partitioning of the two dyes is obtained. In fact within the detection limit of our microscope, which approaches single molecule sensitivity, no fluorescent signal from DiD in the nanotube membrane segment is observed. Interestingly the two dyes were shown to segregate during endocytosis^16^. Prompted by this finding we investigated the nanotube composition using a series of fluorescent staining. The nanotube segment is rich in membrane protein as seen by NHS-Atto647, which forms complexes with protein primary amines. Nanotubes were found negative for clathrin coats evidenced by absence of Transferrin marker and also found negative for Annexin A5 binding. They were positive for staining with cholera subunit B (CT-B) (**Fig. 2e**). These results are consistent with a cylindrical nanotube morphology which was shown to reduce Annexin A5 binding affinity^17^ and nanotubes found in cells for non-clathrin mediated endocytosis marker CT-B^4,18^. Taken together these results show strong partitioning of lipids and protein in the curvature field of the two membrane segments, effectively coupling membrane composition, phase state and curvature^19^. These results are in agreement with prediction from molecular dynamics simulation of lipid partitioning in nanotubes formed from PM lipid mixtures^20^, further corroborating our observations. In fact we could have arrived at a similar conclusion by observing the tGPMV membrane dynamics: Upon osmotic deflation the outer membrane segment starts to fluctuate with optically resolvable amplitudes, indicating a membrane tension in the entropic regime^21^. This implies that the outer membrane is close to its preferred membrane curvature^22^. Taking tGPMV diameter of about 5 μm yields *m_outer_* < 1/5 μm^−1^ for the spontaneous curvature of the outer membrane, while the nanotubes have a spontaneous curvature of *m_tubes_* > 10 μm^−1^. Thus, tGPMVs must exhibit two membrane segments of different composition.

Combining our results we see, that when the plasma membrane is left to relax and equilibrate from cytoskeletal pining, we generate vesicles that exhibit two membrane segments of different composition and membrane curvature. Together with the sharp contrast in lipid order we find that the outer membrane segment and nanotubes exhibit the hallmarks of a liquid-liquid phase separation.

## Membrane curvature and phase separation act to stabilize area reservoirs and lead to super-elastic response

We now aim to find a minimal model for the cell-like elasticity upon aspiration of tGPMVs. The elastic response (**Fig. 2a**) implies a change in tGPMV free energy *F*, which appears as an elastic modulus 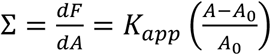. As shown above, nanotubes and outer membrane segment have distinct composition. Membrane tension applied to the outer membrane segment will recruit some of the membrane area stored in nanotubes and the previously (spontaneously) segregated components will have to (re-)mix. Note that we do not observe macroscopic domain formation on the outer membrane segment during an aspiration trajectory. Thus, one contribution to the free energy increase with applied tension will be the mixing of the membrane components to a homogenous phase during aspiration. Considering this effect by direct lipid-lipid interactions, we find in first approximation (see Methods and Materials)

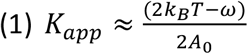

which for typical values of the lipid area *A*_0_ = 0.6*nm*^2^ and lipid-lipid interaction parameter in the order of the thermal energy ω ≈ *k_B_T* (see Methods for details) gives *K_app_* ≈ 3.4 mN/m. This value is close to the measured value of 3.1 +/− 0.2 mN/m. In principle there might be additional contributions to the tGPMV free energy, including scaffolding of proteins ^23^, line tension effects, crosslinking between the nanotubes or changes of the elastic parameters of the membrane itself ^24^. But this minimal model already seems to explain the experimental evidence in the form of a composition change of the outer membrane segment (**Fig. 2d**).

Note that we do not assume that after equilibration with the nanotube segment the outer membrane is not strained, meaning that the mechanical tension should remain close to zero during an aspiration trajectory of a tGPMV. This is consistent with constant tension response measured on plasma membrane spheres by buffering of membrane area by plasma membrane area reservoirs^25^.

## Bottom-up construction of super-elastic vesicles

After we have identified the physics behind super-elasticity in tGPMV, we demonstrate that vesicles with similar size and elastic properties can be created in the lab *de-novo* from synthetic lipids. In synthetic giant unilamellar vesicles (GUVs) nanotubes are generated by membrane spontaneous curvature, which is induced by membrane asymmetry^6^. In a method developed previously by us, membrane asymmetry is induced by asymmetric distribution of charged lipid DOPG between membrane leaflets^26^. Upon osmotic deflation, DOPG:cholesterol GUVs form inward pointing nanotubes, adopting a similar vesicle morphology as tGPMVs (**Fig. 3a**). However, in contrast to tGPMVs, DOPG:cholesterol GUVs exhibit nanotubes *of the same composition* as the outer membrane segment (**Fig. 3b** red trace). Upon application of membrane tension exceeding a small critical tension, nanotubes in DOPG:cholesterol GUVs flow into the outer membrane at *constant tension*. This leads to a droplet-like instability of the GUV in the glass-capillary and no elastic response^13^.

**Figure 3.**
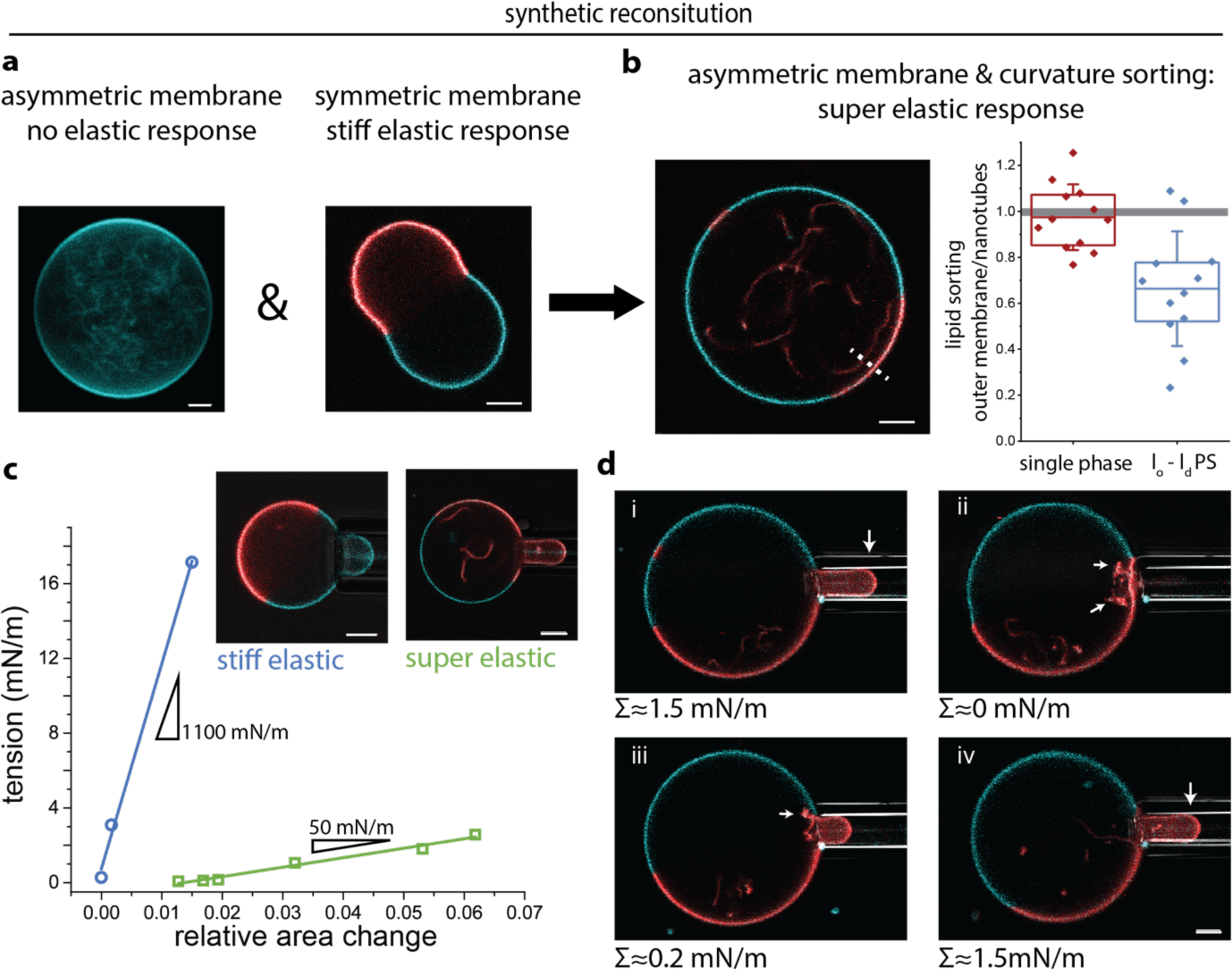
**a)** Tubulated DOPG:cholesterol GUV with asymmetric membrane and symmetric DOPG:SM:cholesterol GUV. Cyan colour shows TopFluor-cholesterol (liquid ordered) and red DiI (liquid disordered). **b)** Asymmetric DOPG:SM:cholesterol GUV. Plot shows normalized membrane composition between outer membrane segment and nanotube (measured along white dashed line) for one phase GUVs (red datapoints) and phase separated (blue datapoints) **c)** Micropipette aspiration data of tubulated and non-tubulated symmetric (blue) and asymmetric (green) DOPG:SM:cholesterol GUV of the same overall composition. **d)** Reversibility of recruitment of nanotubes from phase separated GUV. Time series from i)-iv) where the intermediate aspiration at tension iii) was equilibrated. All scale bars 5 μm.

To create a barrier for the mixing of nanotube and outer membrane material, we introduce sphingomyelin as a third membrane lipid. Sphingomyelin tends to from a liquid ordered phase with cholesterol and to segregate away from DOPG in a liquid-liquid phase separation ^27^. This process is directly visualized by staining with two dyes with partitioning preference for either of the phases (**Fig 3b** blue trace). When DOPG is distributed asymmetrically between leaflets, the liquid-disordered (DOPG-rich) phase forms nanotubes. The dye distribution between the liquid-disordered quasi-planar segment of the outer membrane and the curved nanotube segment reveals that now the composition between the liquid-disordered membrane segments is different. This is consistent with previous reports that sorting of lipids in a curvature field requires direct lipid-lipid interactions and proximity to a demixing transition ^28^. In asymmetric phase separated GUVs, nanotube formation and lipid sorting proceeds spontaneously and is consistent with our hypothesis, nanotube recruitment now proceeds with an apparent super-elasticity. DOPG:SM:chol GUVs appeared elastic and over an order of magnitude softer and about 3 times more deformable compared to the bare symmetric DOPG:SM:chol membrane without any area reservoirs (**Fig 3c**). Note that both types of GUVs have the same overall membrane composition but only exhibit a structural difference, namely in their membrane asymmetry. The deformations in tubulated GUVs were reversible, allowing for recruitment or reforming of nanotubes (**Fig. 3f**).

## Discussion

We found that PM membrane isolated to preserve its native structure, shows similar elastic properties to cells when probed by micropipette aspiration. We have shown that this is due to a lipid nanotube network of distinct composition, exhibiting properties of liquid-liquid phase separation. Plasma membrane extracts do not only exhibit planar phase separated domains ^29^ but are positioned to couple phase state and membrane curvature. While we believe that the high density of the tGPMV nanotube network is formed due to the unphysiological isolation procedure and loss of cytoskeletal pinning and tension, the nanotubes bear some similarities to the CLIC/GEEC nanotubes found *in-vivo:* Similar to our nanotubes, CLIIC nanotubes were found positive for membrane dye FAST-DiI, negative for Transferin maker and of about 40 nm diameter ^4^. Interestingly CLIC/GEEC nanotubes were implicated before in PM tension regulation ^30^. We have shown, that understanding of the main features of nanotube formation stabilized by membrane spontaneous curvature and lipid-lipid interactions enable bottom-up synthetic reconstitution of super-elastic GUVs. Liquid-liquid phase separated membranes were widely studied in their phase behaviours at varying temperature, with small effects of mechanical tension ^31^. By convergence of membrane curvature and liquid-liquid phase separation we obtain responsive synthetic membranes which we anticipate to exhibit rich phase behaviour with varying tension. This provides the ability to sort and reorganize lipids and proteins on plasma membrane mimetics by application of mechanical cues. In this way super-elastic membranes are bound to have a wide applicability in the reconstitution of cellular features requiring membrane remodelling.

Finally, let us mention that super-elasticity is also found in engineered alloys, like Nitinol, and is also based on a phase transition of the host metal^32^. However, tGPMV exhibit threedimensional hysteresis-free remodelling of the nanotubes into outer membrane, demonstrating the remarkable features of biomembrane fluidity.

## Materials and Methods

### GPMV and tGPMV generation and isolation

PM membrane blebbing was induced according to previously published protocols ^10^ with modifications: U2OS cells were cultured in DMEM medium (10 % FBS, 1% Penn-Strep) to 70% confluency, washed 3 times with PBS (phosphate buffered saline), and incubated at 4°C for 10 minutes in PBS (supplemented with 1ul 10mg/mg FAST-DiI or DID in ethanol for experiments shown in Fig. 1b and Fig. 2a – d), washed 3 times in buffer (10 mM HEPES, 150 mM NaCl, 2 mM CaCl2, pH7.4) incubated for 1 hour with 2 mM NEM in buffer. As indicated in the main text in some experiments tGPMV induction was supressed by addition of 25mM Paraformaldehyde (PFA) to the chemical NEM, which instead lead to the formation of ordinary GPMVs. PM membrane blebbing induction with 2mM DTT and 25mM PFA in buffer yielded similar results in terms of GPMVs without reconstitution of a tubular network. Cells were incubated for an hour at 37 °C and GPMV/tGPMV were harvested. Here special precaution was taken to minimize agitation of the flasks. Vesicles were isolated by a single pipetting from a slightly tilted T-75 flasks yielding 1 mL supernatant. This procedure minimized contaminations from cells and cell debris. After additional staining steps (see below) this 1 mL solution was left to stand upright in an Eppendorf tube (1.5 mL volume) to let vesicles sediment by gravity for at least 30 minutes. A 100 μL sample from the bottom of the Eppendorf tube was then further analysed within 12 hours of sample isolation.

### General sample preparation

Generally GPMV and tGPMV were transferred to cleaned (rinsing in ethanol and Millipore water) and bovine serum albumin (BSA) coated coverslip. 100 μL 1 mg/ml BSA was incubated on the coverslip for at least 30 minutes and washed with Millipore water. Between 30 and 50 μL of the tGPMV/GPMV solution were pipetted on the coverslip and sealed with a top cover glass separated by a spacer.

### Staining with water soluble dyes

Water soluble dye and conjugated proteins were prepared according to the manufactures instructions and added to tGPMV and GPMV solution after isolation from the cell culture in the following final concentrations: Alexa Fluor 647 NHS-Ester (Thermo Fisher) at 400 μM, sulforhodamine B Sigma) at 2.5 μM, Cholera toxin subunit B – Alexa Fluor 488 (Invitrogen) at 16.7 nM, Human Transferrin – CF488A (Biotium) at 130 nM, Annexin V – CF594 (Biotium) at 65 nM. Before imaging with the appropriate excitation and emission setting, tGPMV/GPMV were incubated for 20 minutes.

### High resolution imaging

To obtain high-resolution images of the tubular network the diffusive lipid motion was supressed by protein crosslinking. Here the crosslinker PFA was added to tGPMV (isolated with NEM) after isolation from the cell-culture. Importantly crosslinking was only effective if PFA was supplemented with 0.17 mM Triton X 100 (Sigma-Aldrich) to increase membrane permeability. If the membrane was not labelled during plasma membrane isolation, STED dye (ATTO 647N DOPE, Sigma-Aldrich) or FAST-DiI was added to cross-linked tGPMV as 1 μl from a 1mg/ml solution in ethanol to 1 ml isolated tGPMV solution. tGPMV were incubated for an hour at room temperature and imagined.

Image deconvolution was accomplished by the integrated deconvolution plugin (Huygens) in the Leica LAS X. Automatic settings deconvolution setting were applied to slightly oversampled confocal images obtained using a 63×1.2 water immersion lens. STED microscopy was performed on an Abberior Instruments microscope with dye excitation at 640 nm and STED depletion at 775 nm using a pulsed laser source. STED alignment was accomplished using gold beads, by adjusting the focus of the excitation beam into the center of the donut-shaped depletion laser. Corrections for mismatches between the scattering mode and the fluorescence mode were achieved using TetraSpeck beads of four colours. Crimson beads of 28 nm diameter were used to measure the resolution of STED which was found to be about 35 nm.

### FLIM experiments

FLIM experiments were conducted on a Abberior Instruments microscope with pulsed 561 excitation. Fluorescent lifetime trances were analysed using Becker & Hickl SPCM. The data was deconvoluted using the instrument response function (obtained by measurements of DASPI dye emission in methanol). Double-exponential fits were performed and weighted average lifetime reported. Fit was performed on the free parameters of amplitudes, lifetimes and parameters “shift” and “scatter” using the SPCImage software.

### Image quantification

Quantification of confocal images was performed using Fiji/ImageJ v. 2.0. For Fig. 2f, the fluorescent signal in a circular section within the tGPMV (radius 1 μm) and membrane signal (radius 0.1 μm) was averaged and the ratio was reported. For Fig. 3b confocal images were obtained on tubulated GUVs and a line profile (with 1 μm) was drawn between the liquid disordered outer membrane segment and nanotube parallel to the outer membrane section. For each membrane segment the ratio of peak intensity between green and red fluorescent channel (Top flour-Chol and DiI) was calculated. The ratio between the green/red signal in the two membrane segments was then calculated.

### GUV fabrication

GUVs were formed by electroformation from a pre deposited lipid film. As indicated 8:2 DOPG:cholesterol or 3:5:2 DOPG:SM:Cholesterol was deposited as a thin film (4 μL lipid 4 mM solution in chloroform spread on an area of about 1.5 × 3 cm^2^, 2 hours drying at vacuum) on ITO glasses. By application of a low AC-voltage during formation (740 mV_rms_, 10 Hz, 2 hours, 60°C) DOPG was distributed asymmetrically in between the leaflets ^26^. GUV were doped with 0.1 mol% DiD and 0.5 mol% Topflour-Cholesterol. DOPG asymmetry decays over time by passive flip-flops. Nanotubes mostly disappeared or widened to optically resolvable diameters by store at room temperature overnight. This population was then used in the “stiff elastic” experiments from Fig. 3 c.

### Micropipette experiments

Micropipettes were prepared from glass capillaries (World Precision Instruments Inc.) that were pulled using a pipette puller (Sutter Instruments, Novato, CA). Pipette tips were cut using a microforge (Narishige, Tokyo, Japan) to obtain tips with smooth inner and outer diameter. Adhesion of the membrane to the pipette was prevented by incubation of the pipette tips in 1 mg/mL aqueous solution of casein or BSA (Sigma). A new pipette was used for the aspiration of each GUV. After the pipette was inserted into the observation chamber, the zero pressure across the pipette tip was attained and calibrated by watching the flow of small particles within the tip. The aspiration pressure was controlled through adjustments in the height of a reservoir mounted on a linear translational stage (M-531.PD; Physik Instrumente, Germany). This setup allowed the pressure to increase up to 2 kPa with a pressure resolution of 1 mPa. The pressure was changed by displacing the water reservoir at a speed of 0.01 mm/s. The displacement was then stopped for 2-3 mins before any data were recorded. Images were analysed using the ImageJ software. The total vesicle area was directly measured from the membrane contour. The relative area change was calculated as 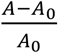 were *A* is the measured vesicle area and *A_0_* is the spherical outer membrane segmented before aspiration.

### Free energy change during GPMV aspiration

We consider a membrane of two segments which are denoted *wc* (weakly curved) and *sc* (strongly curved). Both segments are able to exchange molecules with each other. Further the chemical potential of the molecules in the *sc* segment is approximated to be fixed and in this way acts as a lipid reservoir of constant chemical potential. In the general case the free energy of each membrane segments consist of multiple terms (Helfrich bending energy, line tension terms etc), but in a first step we approximate the free energy of the outer membrane as a binary mixture on a lattice of two equally sized molecules with mole fraction x_1_, x_2_ and total number of molecules N, on a lattice. The free energy of the *wc* segment is then

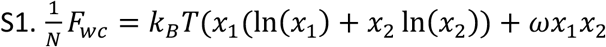

Where *ω* describes the degree of non-ideal mixing between the molecules. Eq. (S1) has an upper critical point 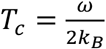 and we consider only temperatures *T* > *T_c_* as the outer wc segment appears as a single homogenous phase (this is strictly only true for tGPMV experiments), which implies 0 < *ω* < 2*k_b_T*. The *wc* segment is assumed to be initially at its energetic minimum, which within the model (S1) implies 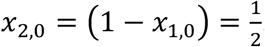. Taylor expansion around x_1,0_ leads to:

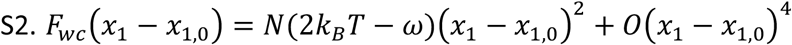

Where constant terms were omitted. Upon aspiration of the outer membrane segment we assume that the outer membrane experiences a transient mechanical tension set by the Laplace pressure on the outer membrane. This mechanical tension relaxes by acquisition of lipid from the *sc* membrane reservoir (nanotubes). We assume that the *sc* reservoir lipids only belong to species *x*_l_. The total number of molecules of species *x*_1_ upon recruitment of Δ*N*_1_ molecules is then 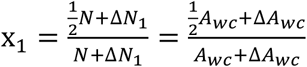 and the total number of molecules in *wc* membrane segment is connected to the area via 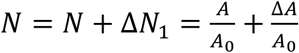 where *A*_0_ is a the molecular area per lipid. Inserting into equation (S2) and taking *A_wc_* ≫ Δ*A_wC_* we find the free energy:

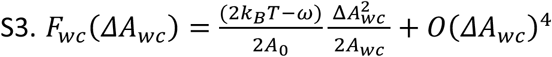

Typical values for *ω* are in the order *ω* ≈ *k_B_T*(23°*C*) ≈ 4.11 10^−21^*J* (ref). This increase in outer membrane free energy leads to the apparent elastic response 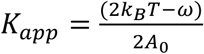.

